# Exploring Mechanisms of Lipid Nanoparticle-Mucus Interactions in Healthy and Cystic Fibrosis Conditions

**DOI:** 10.1101/2024.01.18.575680

**Authors:** Belal Tafech, Mohammad-Reza Rokhforouz, Jerry Leung, Molly MH Sung, Paulo JC Lin, Don D Sin, Daniel Lauster, Stephan Block, Bradley S. Quon, Ying Tam, Pieter Cullis, James J Feng, Sarah Hedtrich

**Affiliations:** Faculty of Pharmaceutical Sciences, University of British Columbia, Vancouver, British Columbia, Canada; Center of Biological Design, Berlin Institute of Health at Charité – Universitätsmedizin Berlin, Germany Department of Infectious Diseases and Respiratory Medicine, Charité – Universitätsmedizin Berlin, Corporate member of Freie Universität Berlin and Humboldt Universität zu Berlin, Germany; Max-Delbrück Center for Molecular Medicine in the Helmholtz Association (MDC), 13125 Berlin, Germany; Department of Chemical and Biological Engineering, University of British Columbia, Vancouver, British Columbia, Canada; Department of Mathematics, University of British Columbia, Vancouver, British Columbia, Canada; Department of Biochemistry and Molecular Biology, University of British Columbia, Vancouver, British Columbia, Canada; Acuitas Therapeutics, Vancouver, British Columbia, Canada; Centre for Heart Lung Innovation, University of British Columbia, Vancouver, British Columbia, Canada; Institute of Pharmacy, Biopharmaceuticals, Freie Universität Berlin, 12169 Berlin, Germany; Institute of Organic Chemistry, Freie Universität Berlin, 14195 Berlin, Germany; Faculty of Medicine, University of British Columbia, Vancouver, British Columbia, Canada Adult Cystic Fibrosis Clinic, St Paul’s Hospital, Vancouver, BC, Canada

**Keywords:** Lipid nanoparticles, Brownian dynamics simulation, cystic fibrosis, particle diffusivity, transmucosal delivery

## Abstract

Mucus forms the first defense line of human lungs, and as such hampers the efficient delivery of therapeutics to the underlying epithelium. This holds particularly true for genetic cargo such as CRISPR-based gene editing tools which cannot readily surmount the mucosal barrier. While lipid nanoparticles (LNPs) emerged as versatile non-viral gene delivery systems that could help overcome the delivery challenge, many knowledge gaps remain, especially for diseased states such as cystic fibrosis (CF).

This study provides fundamental insights into Cas9 mRNA or ribonucleoprotein-loaded LNP-mucus interactions in healthy and diseased states by assessing the impact of the genetic cargo, mucin sialylation, mucin concentration, ionic strength, pH, and polyethylene glycol (PEG) concentration and nature on LNP diffusivity leveraging experimental approaches and Brownian dynamics simulations.

Taken together, this study identifies key mucus and LNP characteristics that are critical to enabling a rational LNP design for transmucosal delivery.

**Graphical Abstract:** 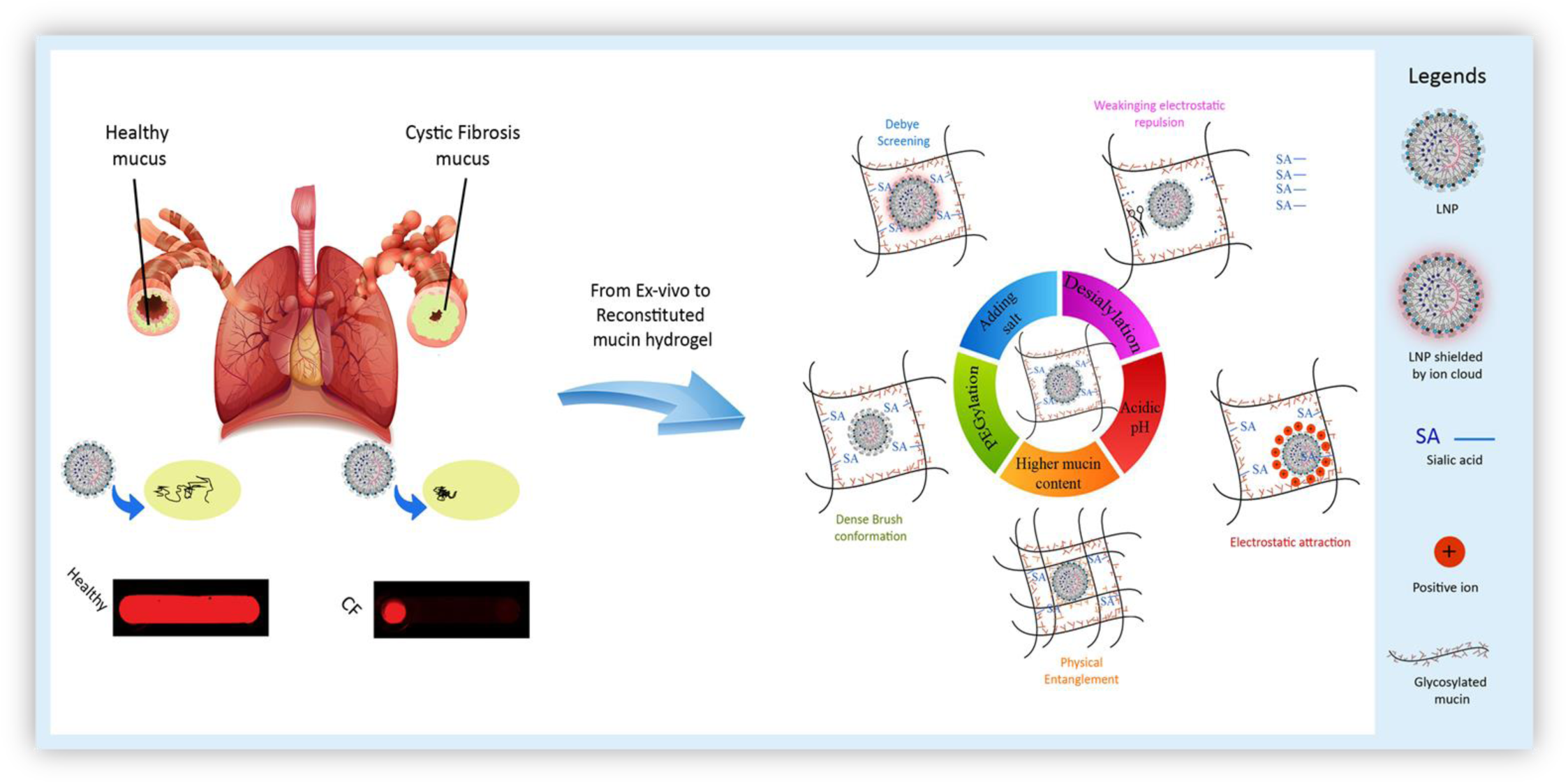

## 1. INTRODUCTION

The mucus layer that lines human organs such as the lungs, the reproductive, as well as gastrointestinal tract, poses a challenging barrier for therapeutics.[1] Successful transmucosal delivery, however, offers great therapeutic opportunities for the treatment of a variety of diseases including monogenic diseases of the respiratory tract such as cystic fibrosis (CF). CF is an autosomal recessive disorder caused by mutations in the cystic fibrosis transmembrane conductance regulator (CFTR) gene and is the most common fatal genetic disease globally.[2] In fact, the median age of survival in highly developed countries such as Canada remains at 33 years only. While significant therapeutic advances have been achieved by the introduction of CFTR modulators [3], there is still no cure for CF. CRISPR-based gene editing, however, now provides us with the tools to potentially enable that.[4] CRISPR/Cas9 together with sophisticated tools such as base or prime editors theoretically can now correct ∼90% of all known disease-causing mutations.[5]

Despite these exciting advances, one main hurdle preventing clinical translations is inefficient transmucosal delivery.[6] Over the past years, lipid nanoparticles (LNPs) emerged as the most promising non-viral delivery system for genetic drugs, yielding great clinical success in delivering siRNA and mRNA.[7,8] LNPs are typically composed of four components: ionizable, helper, and PEG-lipids as well as cholesterol.[9] The ionizable and helper lipids enable efficient cargo encapsulation and facilitate cell uptake and endosomal release. Cholesterol increases LNP stability and promotes membrane fusion during cellular uptake.[10] PEG-lipids decorate the LNP surface and, thus prevent particle aggregation.[10,11] Also, in the context of transmucosal delivery, PEG can minimize mucus-nanoparticle (NP) adhesion and facilitates penetration.[12–15] Despite increased efforts to shed light on NP - mucus interactions, many knowledge gaps remain, especially for diseased mucus.[16]

Normal airway mucus consists of ∼ 97-98% water and solid components such as mucins (∼ 2%), salts (∼ 1%), lipids, DNA, and cellular debris.[17–19] CF mucus is characterized by significantly ∼ 10% higher mucin concentrations, smaller mucus pore sizes (60-200 nm *vs.* 100–500 nm in healthy mucus)[12,20–23], elevated ionic concentrations[24], and lower pH values[25]. This renders CF mucus highly viscous (322 ± 199 Pa-s compared to 10 Pa-s of healthy mucus) ultimately reducing particle diffusion.[12,26–30] In fact, mucus rheology is contingent on the mucin concentration whereas a 2-fold increase in mucin concentrations results in 6-10-fold increased viscosity.[31]

In addition to viscous and steric hindrance, chemical and electrostatic interactions between NPs and mucus are important. For example, the mucin polypeptide is heavily decorated with oligosaccharide chains, constituting 70–80% of the total mucin mass. These oligosaccharide chains are usually terminated by sulfate, sialic acid (SA) or fucose.[32–35] Terminal SA groups are particularly important as they contribute significantly to its net negative charge.[35] At present, the effect of SA on transmucosal delivery is controversially discussed; both suppression and promotion of particle diffusivity have been reported.[36,37]

The ionic strength and acidity of the mucus can also affect electrostatic interactions between NPs and mucin. Salt ions form a double layer around charged NPs and shield their surface charges.[27] The elevated salt concentrations in CF mucus may result in a stronger shielding effect, thus ultimately influencing NP transport. Finally, mucus exhibits tissue-and disease-dependent pH levels.[38] For example, newborns with CF produce moderately acidic (pH 5.2) airway mucus.[39] Since mucin structure and overall charge as well as NP charge are pH sensitive, the mucus pH may affect electrostatic NP-mucus interactions and thus the efficiency of therapeutic delivery.[40,41]

Further, NP surface chemistry can significantly impact their interactions with mucus. A prime example is PEGylation, which enables polymeric NPs ≤ 200 nm to pass through airway mucus[12–14] due to PEG’s amphiphilic nature and neutral charge. While PEG’s role in other NP types has been well studied, little is known about how much LNP diffusivity depends on PEG density.

Being particularly interested in LNP as a carrier for genetic cargo, we investigated the interfacial interactions of LNPs loaded with Cas9 mRNA and ribonucleoprotein (RNP) complexes with mucus hydrogels representative of healthy and CF-like states. More specifically, we investigated the impact of (1) mucin sialylation, (2) ionic strength, (3) pH, and (4) mucin concentration and (5) LNP PEGylation on LNP diffusivity in mucus hydrogels.

We coupled Brownian dynamics (BD) simulations with experimentation to provide mechanistic insights for our observations focusing on steric and electrostatic interactions, aiming to aid the rational design of LNPs yielding efficient transmucosal airway delivery. Finally, we demonstrate that modifications to the nature and density of PEG-lipids may hold the key to significantly advancing mucus diffusivity in human CF mucus. Taken together, our study grants new insights into fundamental mechanisms of LNP-mucus interactions and provides rational design criteria for mucus-penetrating LNPs yielding efficient transmucosal delivery of genetic cargo into the lungs.

## 2. RESULTS & DISCUSSION

### 2.1 Diffusivity of LNP in healthy mucus and CF mucus

While successful LNP-mediated gene delivery to the lungs has been reported in mouse models[15,42–44], the translatability of these results to humans is unclear as murine lungs produce significantly less mucus than humans and have a different anatomical layout rendering gene delivery to mouse lungs much easier. Due to ethical concerns over harvesting lung mucus from healthy individuals, we used porcine lung mucus for our studies, which has comparable physical and chemical properties.[45] To study LNP behavior in diseased states, spontaneously produced mucus samples from CF patients were used.

CRISPR-Cas9-based gene editing now provides us with the tools to potentially cure monogenic lung diseases like CF.[46,47] Yet, the challenge remains that there is still no efficient way of delivering gene editing tools across the highly viscous CF mucus. Gene editing tools can be administered in different formats whereas Cas9 mRNA (complexed with gene-specific sgRNA) and the functional CRISPR-Cas9 ribonucleoprotein complex (RNP) are most commonly used.[48] Both formats offer certain advantages which are extensively discussed elsewhere.[49] However, their impact on the transmucosal diffusivity of non-viral vectors like LNPs has not yet been determined.

Thus, we first assessed the mean diffusivity of unloaded (38 ± 9 nm), Cas9 mRNA-loaded (42 ± 16 nm) and RNP-loaded (279 ± 9 nm) 1.5% PEG-LNPs (**Figure 1**A) in healthy lung mucus. The diffusivity was measured using both multiple particle tracking analysis (MPT) (Figure 1B-C) and a parallel channels method (**Figure 2**A). In the parallel channels, the diffusion of LNPs from one end of the channel to the other was compared by semi-quantification of the fluorescence intensity. It was evident that unloaded LNP and LNP-mRNA had higher diffusivity in mucus, while the larger LNP-RNP diffused less efficiently (Figure 2A). This was confirmed by MPT which recorded the movement and trajectory of ∼266 LNPs on average. Subsequently, LNP displacement and trajectories were analyzed to determine the LNPs diffusion coefficient. The median diffusion coefficient of LNP-mRNA (0.68 µm^2^/s) was twice as high as that of LNP-RNP (0.31 µm^2^/s) (Figure 2B; Supplementary Videos 1 & 2) which is most likely due to the significant size difference.

**Figure 1.**
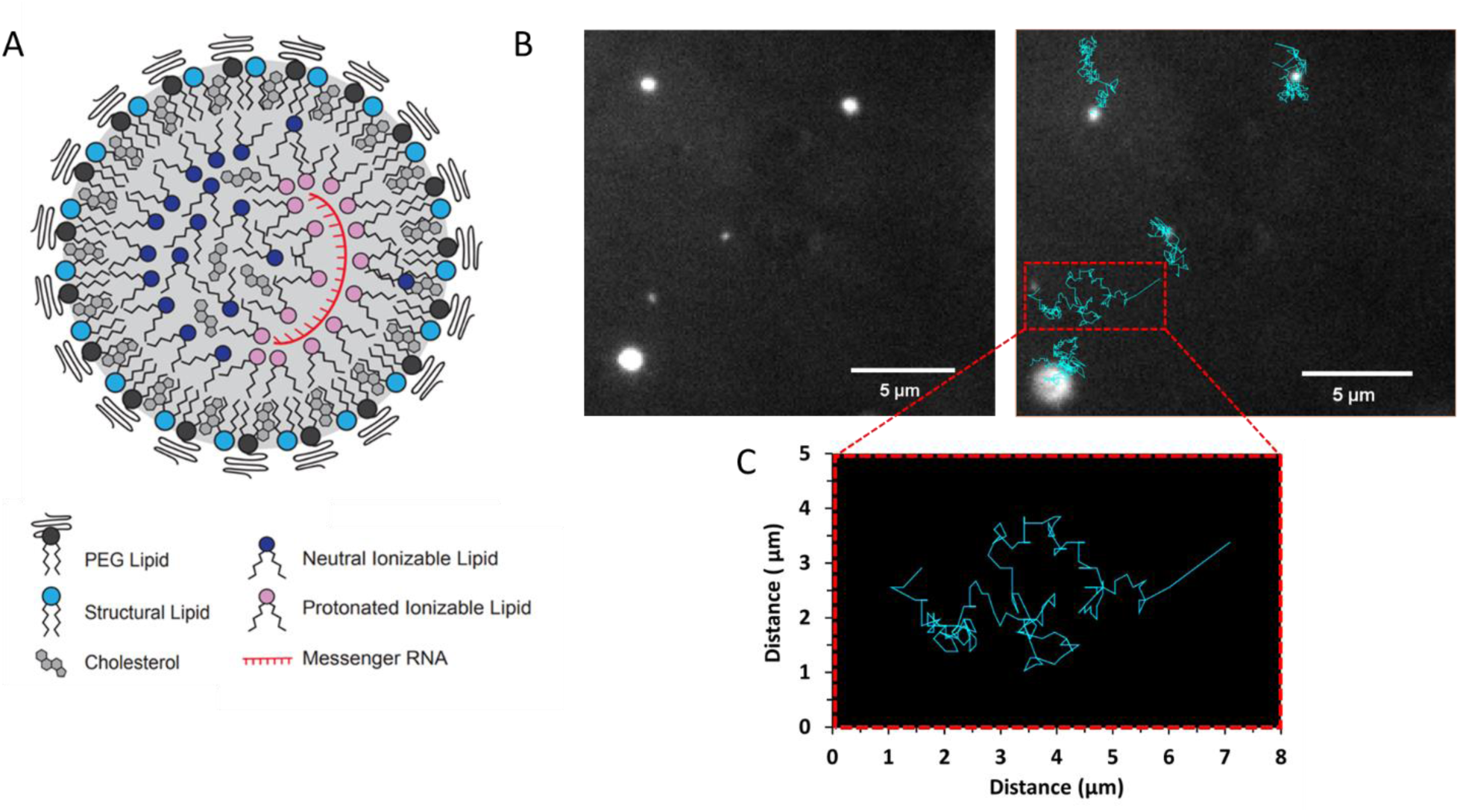
A) Schematic depiction of an LNP encapsulating mRNA (LNP-mRNA). The LNP consists of PEG lipids, helper lipids, ionizable lipids, and cholesterol with the mRNA residing in the core of the lipid complex. B) Schematic depiction of multiple particle tracking analysis (MPT) for determining LNP diffusivity. Top left: LNPs moving in mucus are recorded (266 LNPs on average). Top right: the LNPs are then selected using the spot assistant tool of the NanoTrackJ function in ImageJ and the LNP displacement and trajectories are recorded to determine the diffusion coefficient of each LNP. C) An exemplary zoomed-in view of the trajectory of one LNP.

**Figure 2.**
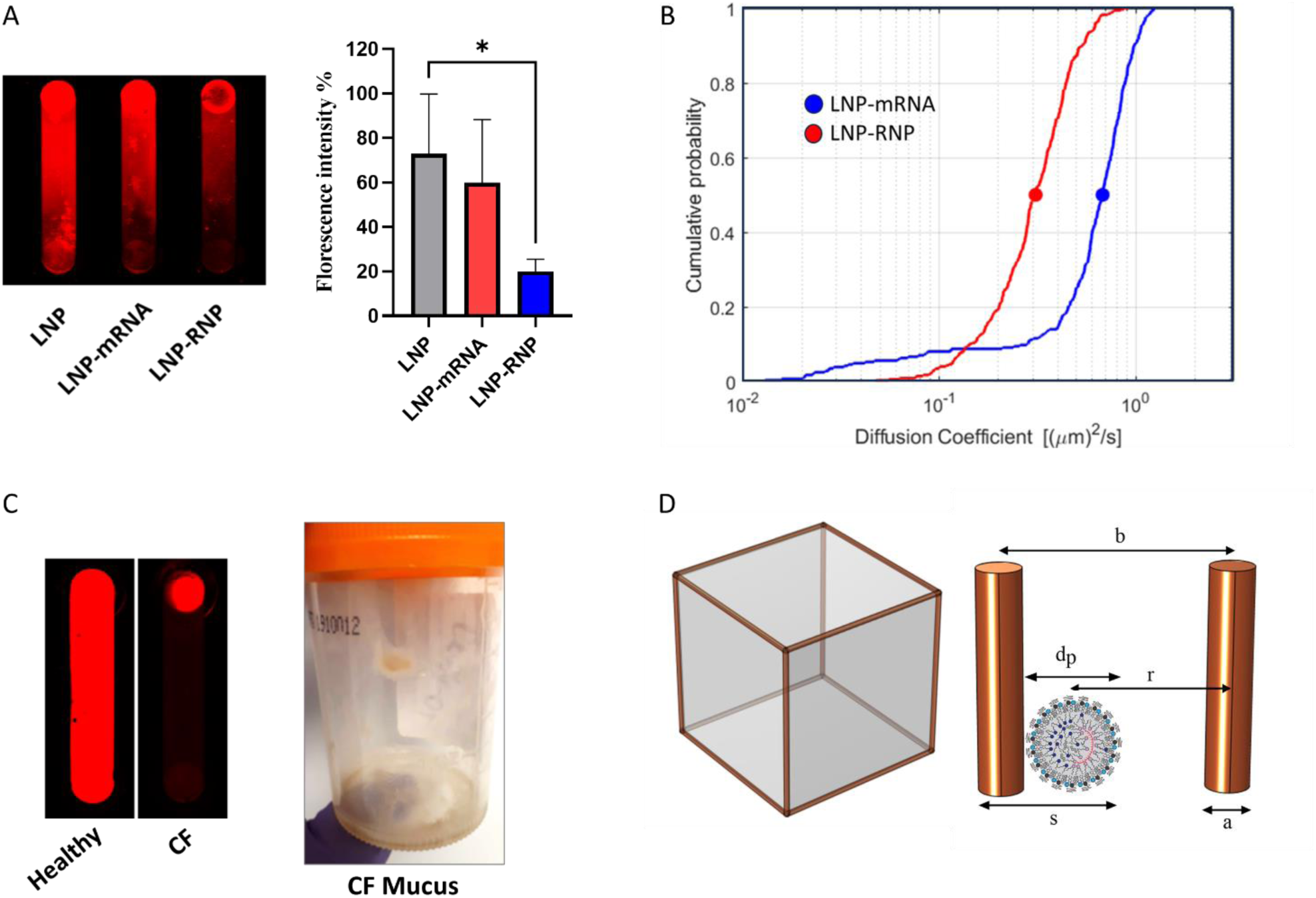
Assessment of LNP diffusivity in healthy and cystic fibrosis (CF) lung mucus via the parallel channel methods and MPT. (A) LNP and LNP-mRNA show greater diffusivity than LNP-RNP in parallel channels filled with healthy lung mucus semi-quantified by relative fluorescence intensities (n=4) *p < 0.05. (B) The median diffusion coefficient of LNP-mRNA is > 2-fold that of LNP-RNP in healthy lung mucus as assessed via MPT. (C) LNP movement in healthy versus CF mucus using the parallel channel method and a representative image of CF patient mucus. (D) Schematic of a cubic lattice used in the Brownian dynamics (BD) simulations to represent the mucin network. The periodic cell has 12 static edges representing mucin chains, and the schematic on the right shows the various lengths of interest: the polymer chain diameter *a*, the particle diameter *d_P_*, the steric diameter *s=a+ d_P_*, the center-to-center distance between the particle and a polymer chain *r*, and the mucus mesh size *b*.

In healthy human lungs, the mucus layer covering the lung epithelium is 10-20 µm thick.[23] In CF patients, however, not only is the mucus more viscous, but the mucociliary clearance is severely impaired which facilitates mucus buildup.[23] The diffusion rate is critical for effective delivery as greater distance must be traveled by the LNPs to reach the underlying epithelium tissue. To emulate that, we repeated the experiment with CF mucus. However, little to no movement of LNPs was recorded (Figure 2C; Supplementary videos 3 & 4) likely due to the significantly higher mucin concentration and smaller pore sizes (60-200 nm)[23]. While we noted distinct patient-to-patient variations in CF sputum structure and rheology[50], the fact that even unloaded LNPs were immobilized indicates that factors other than steric hindrance hamper LNP diffusivity.

### 2.2. Electrostatic interaction between LNPs and mucin hydrogel

To better control for, modify and identify contributing factors to mucus diffusivity such as mucin sialylation, ionic concentration, pH, and mucin concentration, we used reconstituted purified native mucin from bovine submaxillary (BSM) gland for the subsequent experiments since they are well-suited for such investigations.[51] Also, we confirmed the comparability of this mucin hydrogel to normal lung mucus via MPT measurements. In fact, the median diffusivity of LNP-mRNA in 2% BSM mucin in PBS (0.60 µm^2^/s) and in EpiLife medium (0.64 µm^2^/s) match that of LNP-mRNA in healthy porcine lung mucus (0.68 µm^2^/s). Considering the superior diffusivity of mRNA-loaded LNPs in healthy mucus, we focused on mRNA-loaded LNPs for the rest of this study.

#### 2.2.1 Sialic acid cleavage from mucin weakens LNP-mucin electrostatic interaction

Mucin chains are predominantly negatively charged due to sialic acid (SA) residues, whereas NPs can be neutral, negatively or positively charged depending on the surface chemistry.[52] Previous studies explored how SA cleavage affects virus diffusion and binding, yielding controversial results. Kaler et al., observed that more SA increases the diffusivity of Influenza A virus[37], while others showed that SA cleavage aided transmucosal virus movement.[36] To our knowledge, the impact of mucin SA on LNP diffusivity has yet to be studied. Thus, we pre-treated mucin with the SA-cleaving enzyme neuraminidase for generating, what we will refer to for simplicity as, “semi-cleaved” (23% ± 1.3% cleaved SA) and “cleaved” (56% ± 0.9%) mucin samples (Figure 3A). We then tested LNP-mRNA diffusion in these SA-manipulated mucin hydrogels using MPT.

**Figure 3.**
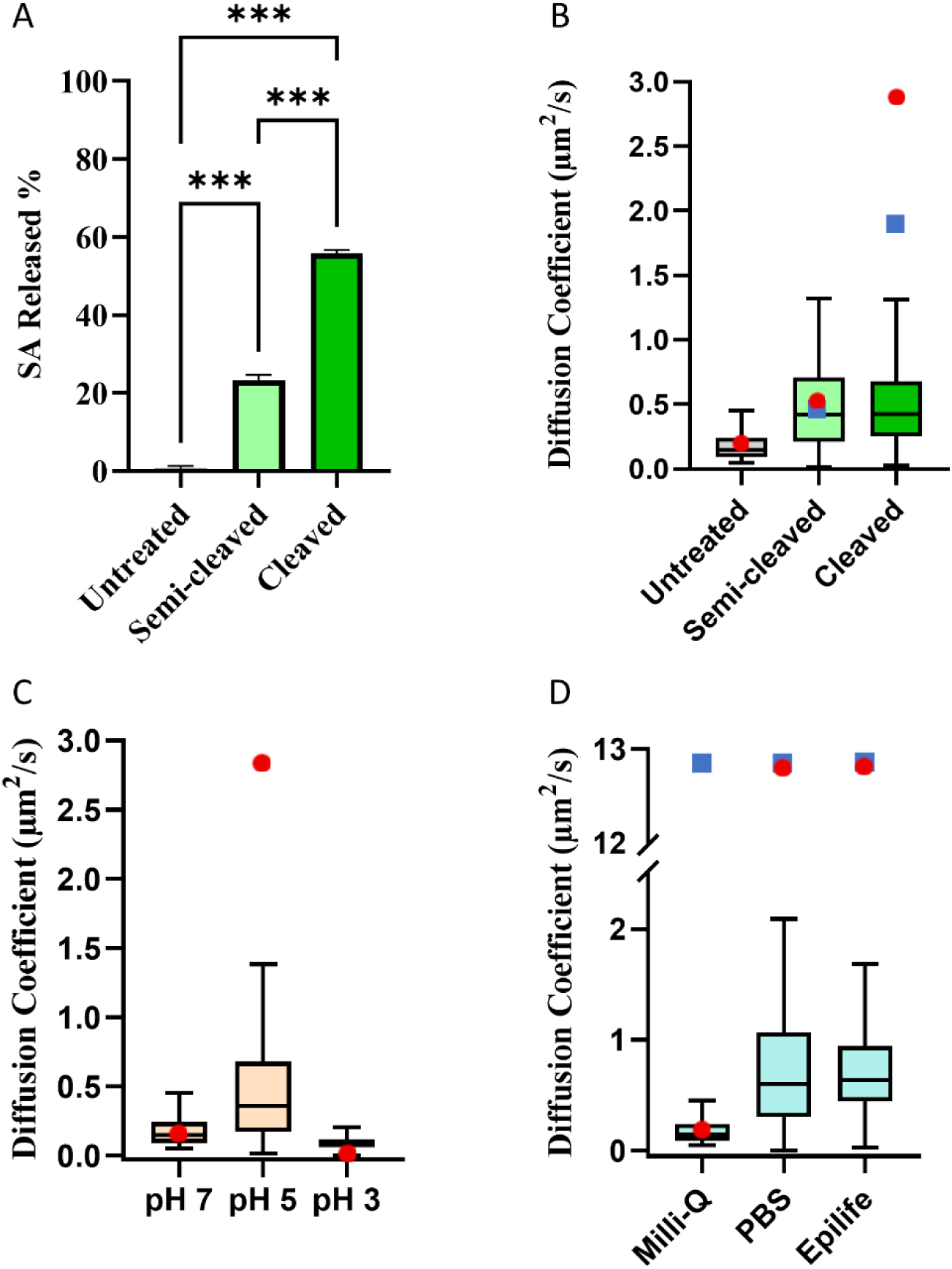
Impact of electrostatic interaction on LNP mucin diffusivity. (A) Percentages of sialic acid (SA) released after 10 min (semi-cleaved) and 4 h (cleaved) of neuraminidase treatment and no treatment (n=3) ***p < 0.001. (B) Diffusion coefficient *D* of LNP-mRNA in the different mucin samples is depicted by Tukey whiskers box plots, where the box marks the 25% and 75% percentiles of the experimental data. Red circles and blue squares show the simulation results based on SA contributing respectively 100% and 85% to the electrostatic interaction. (C) LNP-mRNA diffusion coefficient *D* in 2% mucin solution as a function of solvent pH. Red circles show the simulation results. (D) Variation of the diffusion coefficient *D* in 2% mucin solutions in the three solvents: Milli-Q water, PBS and EpiLife. Blue squares and red circles show simulation results in the absence and presence of the electrostatic interaction, respectively.

In our setup, the median LNP-mRNA diffusivity increased with SA removal (Figure 3B). More precisely, in semi-cleaved mucin hydrogel, the median diffusivity (D = 0.42 μm^2^/s) is 3-fold higher than in the untreated mucin hydrogel (D = 0.15 μm^2^/s). Interestingly, additional SA removal did not further increase the median LNP diffusivity (D = 0.42 μm^2^/s). Since SA is negatively charged, one could ascribe the above trend to the loss of negative charges from the mucin chains, reducing mucin-LNP electrostatic interactions, thus enhancing mobility. But the quick saturation of the median diffusivity with SA cleavage (Figure 3B) is intriguing.

To explore the underlying mechanisms, we employed BD simulations using untreated mucin hydrogel as the baseline (Figure 2D). See Section 3.6 for details of the model and the computational methodology. The LNP diffusion is simulated by solving the Langevin equation subject to Brownian, steric and electrostatic forces. The electrostatic interaction is represented by two parameters: the potential strength *U_e_* and the Debye length *k*. First, we determined these values under a baseline set of experimental conditions (untreated mucin hydrogel), and then varied *U_e_* and *k* according to the experimental modifications (degree of SA removal), with *U_e_* reflecting the mucin and LNP charges and *k* the ionic strength of the medium.

To evaluate the electrostatic potential, we fitted the measured median diffusivity for the baseline case of untreated mucin to get *U_e_/k_B_T* = -80.9. From this baseline, we estimated *U_e_* for the semi-cleaved and cleaved mucin hydrogels in proportion to the amount of charges left on the mucin polymer chains. First, we assessed the diffusivity assuming that the SA domains contributed to 100% of the negative charge on the mucin chains. In reality, however, other charged domains, e.g., sulphate groups, also contribute. However, it is estimated that SA domains constitute ∼85% of the total mucin charges.[53]

Under either assumption (100% or 85% SA contribution), the simulations resulted in diffusivity values closely matching the experimental values in the semi-cleaved group (Figure 3B). However, the simulations predicted higher diffusivities in the cleaved group than what was observed experimentally (Figure 3B). Lowering the charge contribution of SA from 100% to 85% resulted in values closer to experimental diffusivity. Together, the simulations and experimental results may indicate that the SA cleavage effect plateaus as only a fraction of the SA side chains is exposed to the outside and able to interact with LNPs, while the rest are inaccessible (sterically shielded). Therefore, whether these inaccessible SA chains are removed via neuraminidase or not, does not further influence LNP’s diffusivity.

In addition, there are reports that mucin rigidity and rheology are changed when mucin sialylation is altered, suggesting that other physicochemical factors then regulate diffusivity [54,55] although this is controversially discussed.[56,57]

#### 2.2.2 Moderately acidic pH offers optimal LNP diffusion

Human pH varies greatly not only in an organ-specific manner but also within the same organ. For instance, along the gastrointestinal tract, the pH fluctuates between pH 1 and 8.[58,59] The pH of the mucus-containing layer which lines the airway epithelium, in both normal and CF airways, ranges between pH 5.5 - 6.7 in the nasal mucosa, and is ∼ pH 7.0 in the bronchia/lungs.[60,61] Interestingly, neonates with CF reportedly have a more acidic pH (pH 5.2) than non-CF neonates (pH 6.4).[39] Hence, the pH of the airway mucus can vary considerably depending on the airway region, developmental stage and interindividual variations.

We therefore measured LNP diffusivity in mucin hydrogels covering a pH range between pH 3 – pH 7. While pH 3 is physiologically irrelevant for the lungs, spanning a pH range provides valuable mechanistical insights. Especially, since the isoelectric point, defined as the pH at which a molecule has zero net charges, is at pH 2-3 for mucins [41], with higher pH resulting in a negative net surface charge.[40] The surface charges on the mucin polymer chains at different pH values were determined in another study [62] and are shown in **Table 1**. Similarly, the measured Zeta potential of LNPs is also impacted by pH (Table 1), resulting in positive Zeta potentials when the pH is reduced from pH 7 - pH 3.

**Table 1.** Zeta potential and electrostatic interaction of mucin and LNP-mRNA at different pH values. The baseline Ue = -80.9kBT is determined by fitting diffusion coefficient *D* for pH 7. The baseline *U_e_* = - 80.9*k_B_T* is determined by fitting diffusion coefficient *D* for pH 7. The *U_e_* values for pH 5 and 3 are estimated by proportion to the product of the LNP and mucin surface charges.[65]

|  | pH 7 | pH 5 | pH 3 |
| --- | --- | --- | --- |
| Mucin Zeta potential<sup>[62]</sup> | -7.7 mV | - 4.8 mV | - 2.2 mV |
| LNP-mRNA Zeta potential | - $10 \pm 5$ mV | - $7 \pm 2$ mV | + $13 \pm 4$ mV |
| $U_e / k_B T$ | - 80.9 | - 35.1 | + 28.9 |
| Type of interaction | Strong repulsion | Weak repulsion | Weak attraction |

Interestingly, our experimental results showed that the intermediate pH 5 resulted in the highest diffusivity (Figure 3C; Supplementary videos 5-7), indicating a non-monotonic pH effect on LNP-mRNA diffusion in mucin hydrogels. This trend was also captured by our BD simulations, which also provides an explanation to this non-monotonic behavior. Based on the individual LNP and mucin surface charges at the different pH points, the LNP-mucin interaction changes from strong repulsion (pH 7) to weak repulsion (pH 5) and finally to weak attraction (pH 3) (Table 1). Previous computations showed that both electrostatic repulsion and attraction can hinder NP diffusion in mucus.[27] When plotted against *U_e_*, therefore, *D* presents a maximum near *U_e_* = 0 (i.e., neutral). As *U_e_* takes on larger negative (attractive) or positive (repulsive) magnitudes, *D* is suppressed. Furthermore, electrostatic attraction is much more effective in trapping NPs than electrostatic repulsion at the same magnitude of *U_e_*. For pH 3, in particular, electrostatic attraction traps the LNPs near the corners of the cubic lattice and suppresses their diffusion greatly.

Although the trend has been predicted by an earlier computation[27], ours appears to be the first experimental demonstration of this effect. The non-monotonic variation of LNP diffusivity with pH disagrees with the only prior experimental data in the literature.[63] Lieleg et al.[63] tested the behavior of much larger PEGylated polystyrene particles (> 1 µm) in gastric mucins and reported a monotonic decrease of *D* with decreasing pH. There is evidence that larger particles may modify or even break the local mucin network, which may account for this discrepancy.[64] Using functionalized silica NPs, Guo et al.[38] reported ligand-specific reactions to pH changes. Their observations that mucus acidity induces positive charges on NP surfaces, and thus tends to immobilize NPs by electrostatic entrapment agree with our results.

#### 2.2.3 Ions screen LNP-mucin electrostatic interaction

Normal lung mucus contains roughly 1% salts such as sodium chloride[66], whereas CF mucus tends to contain a 30% higher salt concentration.[24,67] CF patients also nebulize different percentages of hypertonic saline to hydrate the mucus which can further increase mucus salinity.[68] Salts dissociate into ions and form a double layer around charged NPs in what is known as the Debye screening effect, which ultimately decreases the electrostatic interaction range, i.e., the Debye length.[27] This suggests that mucosal ionic strength may have practical implications regarding LNPs transmucosal delivery.

To quantify the effect of ionic strength, we tested LNP-mRNA diffusivity in 2% mucin hydrogel made with phosphate-buffered saline (PBS) as the solvent. PBS has an ionic strength of approximately 160 mM, whereas the deionized Milli-Q water has an ionic strength of ∼0.01 mM. The results showed that diffusivity increases with the ionic content, from 0.15 µm^2^/s in Milli-Q water to 0.60 µm^2^/s in PBS (Figure 3D). To test whether the ionic strength impact on LNP diffusivity applies in a more complex cell culture medium, the measurement was repeated in EpiLife, an epithelial cell culture medium (160 mM). The diffusivity of LNP-mRNA was 0.64 µm^2^/s, which is almost identical to that of PBS (Figure 3D). These results indicate that ionic strength is an important diffusivity determining factor even in more complex solvents such as EpiLife which contains additional factors such as amino acids and proteins.

To further investigate the ionic strength-diffusivity relationship, we employed BD simulations. To probe the impact of solvent salt in our BD simulations, we assume that the electrostatic potential remains at the baseline value for the three solutions: *U_e_*/*k_B_T* = -80.9. Thus, the solvent affects the NP diffusivity only through electrostatic screening, represented by the Debye length *k*. For Milli-Q water, with negligibly low ionic content, we adopt a large Debye length *k =* 20 nm from prior literature.[69] PBS and EpiLife have a higher ionic strength ∼ 160 mM[70] which corresponds to a Debye length of *k =* 0.75 nm (see Eq. (4) in the Supplementary Materials).

The simulations reproduce the trend that *D* increases with ionic strength (Figure 3D). As expected, since PBS and EpiLife have the same Debye length, they show identical behavior. However, the computed *D* is much higher than the measured values for PBS and EpiLife. This implies that in reality, the ions in these solvents do screen the LNP-mucin electrostatic repulsion to raise D, but not nearly to the same degree as expected theoretically from the ionic concentration of these solvents. To probe this discrepancy further, we conducted simulations with electrostatic repulsion turned off (Figure 3D). The differences between the blue and red circles show the significance of the electrostatic interaction. In Milli-Q water, electrostatic repulsion is effectively unscreened and plays a major role in suppressing D. In PBS and EpiLife, on the other hand, the model predicts nearly complete screening by free ions at 160 mM rendering electrostatic repulsion negligible. This did not happen in the experiment.

Interestingly, the effectiveness of double-layer screening has been a long-standing but rarely discussed puzzle in the literature. Previous computations[27,71,72] have shown that according to the double-layer theory, ionic concentrations ≥100 mM should shield the electrostatic interaction completely, in agreement with our own computations. However, our own and other experimental results [73,74] indicate that, at this level of ionic strength, electrostatic interaction continues to play a considerable role in suppressing LNP transport through mucus. One possible explanation for the discrepancy is that the mucin structure may change under different ionic strengths. For instance, the flexible non-glycosylated regions of the mucin backbone take on more compact conformations at higher salt concentrations[75], which may lead to higher intrinsic viscosity slowing down NP diffusion. Such structural changes have not been accounted for in the simulations.

### 2.3. Surface PEGylation improves LNP diffusion

Surface PEGylation can shield the particle core from adhesive interactions with mucus owing to PEG’s amphiphilic nature and neutral charge, allowing the particle to move more freely.[76] This has been demonstrated by decorating different types of NPs up to 200 nm with PEG.[12–14] For LNP, the PEG-lipid is one of usually four lipid components. So far, the discussion of its role has mostly focused on its effect on LNP structure and cellular uptake, but its impact on LNP’s mucus diffusivity is scarcely studied.

To probe the impact of PEG concentration on LNP diffusion, we first varied the molar concentration of the commonly used DMG-PEG 2000 from 1% to 5%. According to prior observations, increased PEG concentrations lead to reduced LNP size *d*_p_ [42,77] and reduced negative charges on the particles (Figure 4A-B).[76] At low PEG concentrations, LNPs fuse into larger particles, whereas denser PEG on the LNP surface inhibits such fusion events yielding smaller particle sizes.[78] The amount of surface charge |ζ| also declines slightly with increasing PEG coverage with the exception of 5% PEG, which could be due to more mRNA associating with the surface, due to the smaller *d_p_* and thus limited space for mRNA encapsulation.

**Figure 4.**
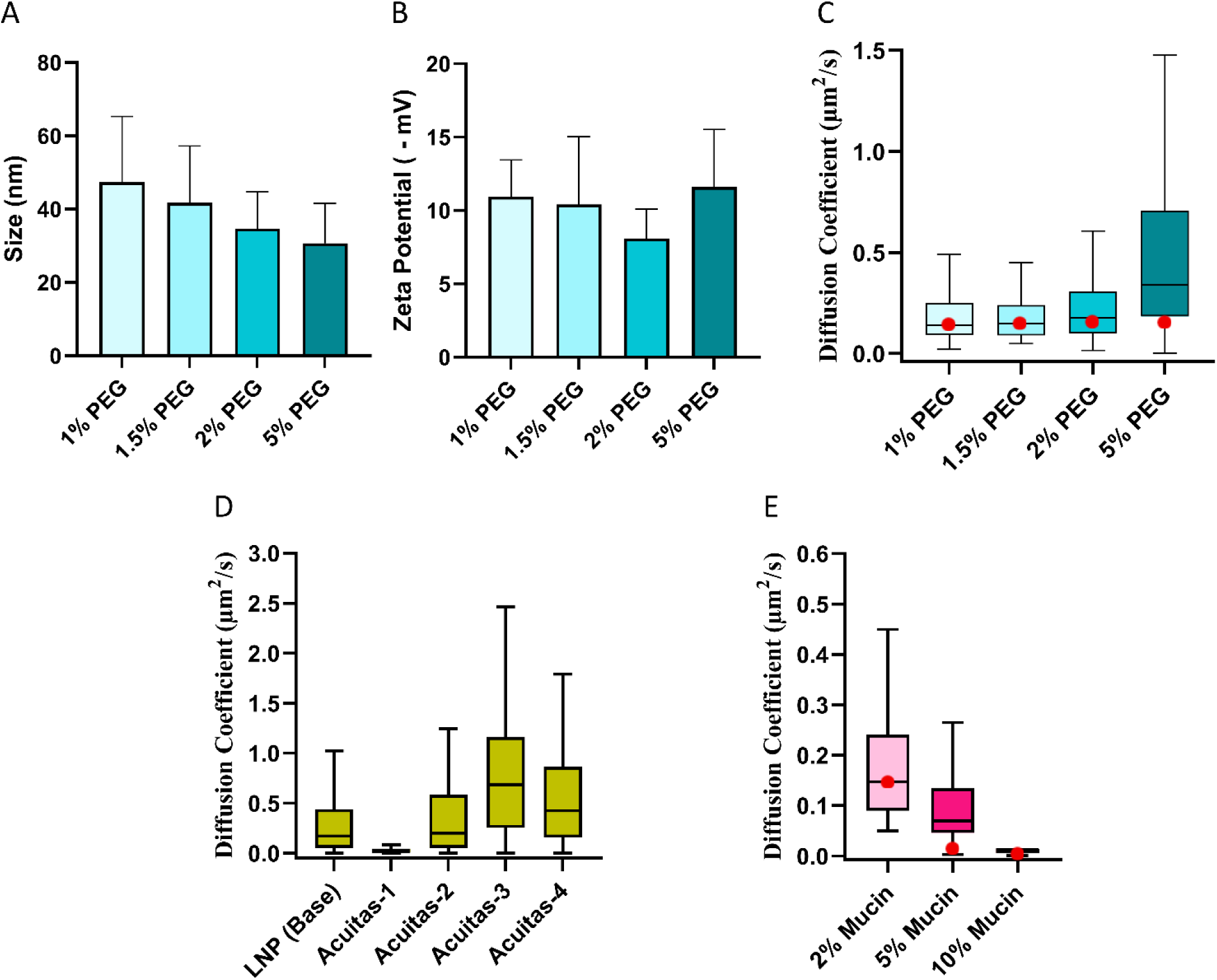
(A) Particle sizes and (B) Zeta potential (ZP) of LNP-mRNA containing 1, 1.5, 2 and 5% PEG. (C) Diffusion coefficient *D* of LNP-mRNA with different PEG concentrations in 2% mucin in Milli-Q water. red circles are diffusivities calculated by numerical simulations. (D) Diffusion coefficient *D* of Acuitas LNPs (1-4) and LNP-mRNA (control, containing 1.5% PEG-2000) in human CF mucus. (E) Diffusion coefficient *D* of LNP-mRNA (with 1.5% PEG) in different concentrations of mucin (2%, 5% and 10%) in Milli-Q water. Red circles are diffusivities calculated by numerical simulations for mucus with homogeneous pore size.

In mucin hydrogels, we observed a gradually increasing diffusivity *D* for increasing PEG coverage (Figure 4C) which is due to both smaller particle size and diminished electrostatic repulsion[42]. BD simulations captured the same trend, adopting the *d_p_* and ζ parameters for each PEG concentration.

Quantitatively, however, the computation predicts a weaker increase of *D* with PEG concentration. This may be due to other changes brought on by PEGylation that are not accounted for by the theory. For example, Xu et al.[79] and Guo et al.[38] observed a transition from mushroom to brush conformation at 5% PEG, which enhances particle diffusivity which is consistent with our experimental data.

It has been reported that increasing PEG density > 3% may reduce cellular uptake[80] due to reduced adsorption of apolipoprotein E (ApoE).[81] This effect, however, was mainly observed for hepatocytes. There have also been concerns regarding the decrease in payload encapsulation capacity of LNPs with high PEG content. For instance, a recent study showed that LNPs with ≥ 5.5% PEG 2000 show decreased payload encapsulation.[42] Thus, while increasing PEG density in LNPs favors transmucosal transport, further research is needed to maintain high payload encapsulation efficiency.

#### 2.3.1. PEG modifications can aid LNPs diffusion in CF mucus

Building onto the potential mucus-enhancing effect of PEG, we investigated if LNP mucus diffusivity in highly viscous CF mucus could be enhanced through PEG modifications. Therefore, we generated four LNPs encapsulating mRNA which varied in the PEG steric barrier (Acuitas 1-4) for testing in human CF mucus. These LNPs differed in either the nature and/or the amount of PEG steric barrier lipid to provide a range of hydrophilic shielding ability. Acuitas-1 and Acuitas-2 contained a single PEG species, whereas Acuitas-3 and Acuitas-4 contained a mixture of PEG species. Neither the amount nor the specific PEG species of these LNPs appeared to have meaningful impact encapsulation efficiency (93-97% determined by Ribogreen); however, the data shows that particles with lower levels of PEG content tended towards a more uniform size as indicated by the polydispersity index (PDI) (**Table 2**). Remarkably, our results showed that LNPs containing a mixture of PEG species had improved diffusion in CF mucus. Specifically, the median diffusivity of Acuitas-3 was 0.69 µm^2^/s which is 4-fold higher than the diffusion rate of our baseline LNP-mRNA formulation (0.17 µm^2^/s) in CF mucus (Figure 4D), equivalent to that of the baseline LNP-mRNA in healthy lung mucus (Figure 2B; 0.68 µm^2^/s). Similarly, Acuitas-4 showed ∼2.5-fold greater diffusivity (0.42 µm^2^/s) than the base LNP-mRNA showcasing PEG and its modifications as an important parameter to facilitate mucus diffusion of LNP in diseased states (Supplementary videos 8 & 9).

**Table 2.** Encapsulation efficiency %, size, and polydispersity index (PDI) of Acuitas LNPs.

| Formulation Name | Encapsulation Efficiency % | Size (nm) | PDI |
| --- | --- | --- | --- |
| Acuitas-1 | 97% | 52 | 0.120 |
| Acuitas-2 | 93% | 40 | 0.144 |
| Acuitas-3 | 97% | 43 | 0.144 |
| Acuitas-4 | 93% | 46 | 0.160 |

### 2.4 Higher mucin concentration hinders LNP diffusion

Mucin concentration varies widely between healthy and disease states. Whereas healthy airway mucus contains ∼2% mucins[17–19], the mucin concentrations in CF mucus range between ∼5% - 10%. This results in smaller pores in CF mucus (60-200 nm vs. 100-500 nm in healthy mucus) which can sterically hinders particle diffusion.[12,20–23] In our experiments, the 2% mucin hydrogel represents healthy mucus while 5% and 10% emulates the mucus found in mild and severe CF cases.[21,22] To systematically assess the impact of mucin concentration on LNP mucus diffusion, we measured LNP-mRNA diffusivity in 2%, 5%, and 10% mucin hydrogels (Figure 4E). Recall that in our BD modeling, the mucin concentration is reflected by the mesh size *b* in the unit cubic cell (Figure 2D), which can be determined from the mucin concentration.

As expected and in line with prior studies[63], the experimental data show that the diffusivity decreases monotonically with increasing mucin concentration (Figure 4E). Consistently, the BD simulation predicts a monotonic behavior, with a sharp decrease in *D* from *C* = 2% to 5%, followed by a slight decrease at *C* = 10%. Overall, these results indicate that the elevated mucin concentration associated with severity of the CF disease can negatively affect therapeutic transmucosal delivery.

## CONCLUSIONS

Lipid nanoparticles (LNPs) are the most advanced non-viral delivery vectors for genetic cargo to date. A strong interest lies in the delivery of genetic material such as mRNA-based CRISPR-systems to the lungs as this opens a new therapeutic avenue for a plethora of diseases. LNPs face a restrictive mucus barrier in the lung [23] which significantly hampers particle penetration. Also, given the short turnover time of lung mucus (10-20 min), the LNPs need to traverse the mucus barrier rapidly to reach the epithelium. The goal of this study was to close critical knowledge gaps in our understanding of LNP-mucus interactions and provide new insights that may aid designing optimized LNP systems.

A critical aspect of mucosal diffusivity is always the particle size. In our experiments, mRNA-loaded LNP showed a much greater diffusivity (0.68 µm^2^/s) than RNP-loaded LNP (0.31 µm^2^/s) in healthy mucus. As both had the same LNP composition, the difference was most likely due to LNP-mRNA being smaller in size. However, almost all tested LNP showed poor to no diffusivity in CF mucus, indicative of the impact of mucus properties such as the pore size and electric charges as determinants for LNP diffusivity in mucus.

To elucidate these effects, we demonstrated the influence of sialylation on LNP diffusivity and that a moderately acidic lung pH (such as in CF neonates) provides optimal conditions for high LNP diffusivity, whereas a neutral pH as it has been reported in CF adults may hamper it. Interestingly, the ionic concentration of the mucin tends to shield electrostatic interactions between mucin and LNP, thus, increasing their diffusivity. In addition, our data show that LNP surface PEGylation increases the diffusivity monotonically with 5% PEG concentration resulting in the highest diffusivity rates. Most interestingly, we demonstrate that using a mixture of PEG species rather than a single PEG species may be a powerful lever to fine-tune LNP diffusivity which is especially important for diseased conditions.

Our findings may guide optimized LNP design and provide insights into which mucosal factors could potentially be temporarily modified to enable transmucosal LNP penetration. This appears clinically highly relevant, especially in light of the recent failure of a clinical trial focused on topical LNP-mediated CFTR mRNA delivery into the lungs. Here, although initial data seemed promising, no improved lung functions were observed in CF patients.[82–84] Transmucosal delivery problems appear to be a major factor.

While our study closes important knowledge gaps, certain limitations remain. For example, in contrast to the *in vivo* behavior of mucus, our experimental and in silico setups were simplified. Also, the experimental setup does not account for the complexities of *in vivo* mucus such as the pore size distribution and spatial heterogeneities.[85] Further, mucin chains are rather flexible and undergo Brownian fluctuations *in vivo*. As such, the polymer network can deform locally to facilitate the passage of larger particles. The electric charges derive from diverse chemical groups along the mucin chains[86] whereas we assumed a uniformly distributed charge density. Similarly, the LNPs have an inhomogeneous charge distribution with ionizable, structural, and PEG lipids located on the outer surface.[42] Thus, our simplified interaction potential (see Eq. (6) in the Supplementary Materials) might not have captured all the subtleties of electrostatic LNP-mucus interactions. Finally, pH-dependent structural changes in the crosslinked mucin network[87,88] were not considered in this work.

## 3. METHODS

### 3.1 Isolation of native physiological and CF patient-derived mucus

Lungs with intact trachea of sacrificed healthy pigs were provided by the Jack Bell Research Centre as well as UBC Centre for Comparative Medicine in Vancouver. To maintain the native properties of the mucus, the lungs were collected and kept on ice immediately after the pigs were sacrificed and then transported (∼30 minutes) to the lab for mucus isolation. While on ice, the trachea and lung tissue were opened, and the mucus was then gently scraped off, to be used immediately or stored at -20 °C until further usage.

For mucus from CF patients, clinicians from Providence Health Care (Vancouver, BC) facilitated the collection of mucus samples spontaneously expectorated produced by CF patients at St. Paul’s Hospital, Vancouver, BC (Ethics approval number: H20-03198). The CF mucus samples were collected from six CF patients (3 females and 3 males). Out of the six patients, five received hypertonic saline, salbutamol and dornase alfa while only one received nebulized colistin and/or budesonide/formoterol. The forced expiratory volume (FEV1) of the patients ranged from 1.65 to 2.89, with the average FEV1 being 2.18. The average age of the CF patients was 35.2 years (Range: 28 - 43).

### 3.2 Preparation of mucin hydrogels

We first dissolved 20 mg of native mucin from bovine submaxillary gland (Sigma-Aldrich, Saint Louis, MO, USA) in acetate buffer 0.05M (pH 5) at 37 °C for 30 minutes. This mixture was then pipetted onto 10 kDA cut-off Pierce Protein Concentrators PES columns (Thermo Fisher Scientific, Burlington, ON, Canada), centrifuged at 15,000 g for 15 minutes, and washed with Milli-Q water, with the centrifugation-washing cycle being repeated twice. Then, the mucin-containing supernatant was resuspended at 20 mg/mL to obtain 2% mucin in Milli-Q water, phosphate-buffered saline (PBS) or EpiLife medium (GIBCO, Grand Island, NY, USA). Similarly, to obtain 5%, and 10% mucin in Milli-Q water, the mucin-containing supernatant was resuspended at 50 and 100 mg/mL, respectively.

### 3.3 Sialic acid (SA) cleavage

One gram of mucin glycoprotein from bovine submaxillary gland was treated with 1U neuraminidase from Vibrio cholerae (Sigma-Aldrich, Saint Louis, MO, USA) in sodium acetate buffer 0.05 M (pH 5) at 37 °C for 10 minutes or 4 hours. Then, DANA (2,3-didehydro-2-deoxy-N-acetylneuraminic acid) (Sigma-Aldrich, Saint Louis, MO, USA) was added at a final concentration of 17.2 μM to terminate the sialidase reaction.[89,90] Mucin samples undergoing the same treatments but without the addition of neuraminidase served as the control. Cleaved SA were filtered out from the mucin solution using the 10 kDA cut-off Pierce Protein Concentrators PES columns.[91] The mucin retentate was subsequently resuspended in Milli-Q water yielding 2% mucin hydrogels.[22]

The filtrate containing the cleaved SA was collected to quantify the amount of cleaved SA in the 10-minute and 4-hour treatment conditions. Here, the Sialic Acid (NANA) Assay Kit (Abcam, Toronto, ON, Canada) was used according to the manufacturer’s instructions. In brief, 50 μL of filtrate samples were mixed with 50 μL reaction mix, which were then incubated at room temperature for 30 minutes. Subsequently, the absorbances at wavelength 570 nm were measured using the μQuant MQX200 (Biotek Instruments, Winooski, Vt., USA) microplate reader. To approximate the percentage of released SA, the amount of SA in nmol was divided by the total SA expected in intact mucin as per the supplier’s specification. The 10-minute neuraminidase treatment resulted in 23% ± 1.3% cleaved SA (semi-cleaved), while the 4-hour treatment resulted in 56% ± 0.9% cleaved SA (cleaved).

### 3.4 Lipid nanoparticle (LNP) preparation and characterization

LNPs were prepared using the bench-top mixing method as previously described.[92] Briefly, the relevant lipids were mixed in ethanol (10 mM) with an aqueous phase (25mM sodium acetate buffer pH 4.0) using a T-junction.[93] The produced suspension was then dialyzed overnight against 1,000x volume of the same buffer to remove the ethanol. Next, the vesicles were removed and concentrated using an Amicon centrifugal unit (10k MWCO; Millipore Sigma, Burlington, MA). The final lipid concentration was measured using the Total Cholesterol Assay kit (Wako Chemicals, Richmond, VA, USA). Each LNP formulation contains the following: ionizable cationic lipid (MC3), phospholipid (DOPE), cholesterol, and PEG-lipid (DMG-PEG 2000) at 50/10/38.3/1.5 mol% respectively (For LNP with 1.5% PEG). To vary PEG-lipid content in the LNP, cholesterol was adjusted accordingly. The fluorescent labels, DiI-C18 (Invitrogen, Carlsbad, CA) were included at 0.2 mol%.

To allow for a direct comparison between mRNA and RNP-loaded LNPs, we used benchtop mixing since the ethanol in conventional microfluidic mixing approaches would denaturate the RNP. Here, a 1:1 mixture of CRISPR-Cas9 mRNA (TriLink, San Diego, CA, USA) to sgRNA (IDT, Toronto, ON, Canada) was prepared. Then, 1 μg of total mRNA (Cas9 mRNA + sgRNA) was added to 0.034 μmol of LNP diluted in 25μM sodium acetate buffer at pH 4, followed by 10 min incubation at room temperature and subsequent dilution of the pH4 buffer with cell culture media to achieve the necessary pH neutralization to complete particle formation.[93] The initial slightly acidic conditions (pH 4) confers positive charge on ionizable lipids and thus the negatively charged nucleic acid can be effectively encapsulated.[94] Benchtop mixing and incubation of LNP with RNA at pH 4 for 10 minutes at room temperature, followed by neutralization with media has been previously shown to be an effective method for preparing RNA encapsulated LNPs even in the absence specialized mixers.[92]

RNP was prepared by mixing sgRNA with the Cas9 protein (IDT, Toronto, ON, Canada) at 1:1 molar ratio in IDTE pH 7.5 buffer (IDT, Toronto, ON, Canada), followed by incubation at room temperature for 10 minutes. Then 1 nmol of this RNP complex was added to 50 nmol of LNP diluted in 25μM sodium acetate buffer at pH 4 and processed as mentioned above.

The LNP sizes and zeta potentials (ζ) were measured using a Zetasizer Nano ZS system (Malvern Instruments Ltd., Malvern, UK) equipped with the Zetasizer software version 7.13 according to standard procedures (Table S1). The LNP sizes were further confirmed via cryogenic electron microscopy (EM) as shown in Fig. S1.

Acuitas LNP-mRNA formulations (Acuitas 1-4) were manufactured using a self-assembling process as previously described.[95] An ethanolic lipid mixture of ionizable cationic lipid, cholesterol, distearoylphosphatidylcholine, polyethylene glycol lipids, and fluorescent lipophilic dye DiI were mixed with a buffered aqueous solution containing mRNA under acidic conditions. The LNP composition is described under US patent WO 2018/081480A. LNP characterization was conducted at Acuitas Therapeutics (Vancouver, BC, Canada). Particle sizes (between 40-60 nm) and polydispersity (<0.200) were determined using dynamic light scattering using a Malvern Zetasizer NanoZS (Malvern Instruments Ltd, Malvern, UK) and encapsulation was determined by ribogreen assay (>90%).

### 3.5 Characterization of mucus-LNP interaction

#### Multiple Particle Tracking (MPT)

First, NPs were added to mucus/mucin hydrogel in an Ibidi µ-Slide VI-0.4 (Ibidi, Gräfelfing, Germany) at a final volume fraction ≤0.2 %. Then, using the PerkinElmer VoX Spinning Disk fluorescence microscope (PerkinElmer, Woodbridge, ON, Canada) equipped with 100X objective, a high-speed Hamamatsu 9100-02 CCD camera and a thermoplate heated to 37 °C, we recorded the live movement of the NPs in mucus/mucin hydrogel/solvent. The frame rate was set to 10 FPS (except for LNP-mRNA vs LNP-RNP MPT: 50 FPS) and the duration of each video was 10 seconds. On average 266 NPs were used to determine the median diffusivity in each condition.

The NPs’ movements in each captured video were analyzed using the NanoTrackJ plugin in ImageJ software (Version 1.53k) (Figure 1B). First, the spot assistant tool was used to specify an appropriate tolerance value and mean filter size to maximize selection of visible NPs while minimizing noise selection. The NanoTrackJ function then followed the trajectory of each NP, and measured the displacements of a diffusing NP. The NanoTrackJ tool was set to include NPs with a minimum of 10 steps per track. The covariance estimator was then used to estimate the diffusion coefficient of each NP based on the measured displacements. Compared to other methods, the covariance method is an unbiased estimator of diffusion under different experimental conditions.[96]

#### Parallel channels method

The channels of an Ibidi µ-Slide (VI-0.4) are filled with 40 uL of mucus/or mucin hydrogel followed by the addition of the fluorescently labeled LNPs. A Sapphire Biomolecular Imager (Azure biosystems, Dublin, CA, USA) was used to capture the LNP diffusion after 1 hour by semi-quantifying the fluorescence intensity of each channel.

### 3.6 Brownian dynamics simulations

To simulate LNP diffusion through mucus, we model the crosslinked mucin chains as rigid edges of a periodic cubic lattice[27], with the lattice size *b* corresponding to the average mesh size of the mucus. This mesh size can be estimated from the mucin concentration *C* from the following formula[27]:

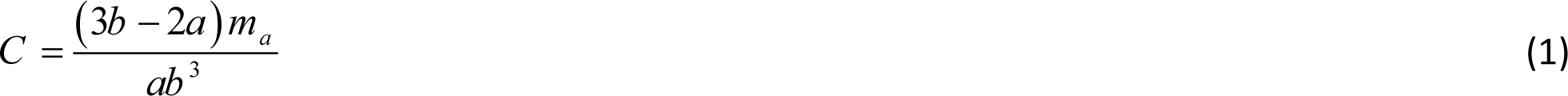

where *a* = 5 nm is the diameter of the mucin chain[32], and m_a_ = 0.3 MDa is the mucin monomer mass.[97] The Brownian diffusion of NPs can thus be tracked within a unit cell (Figure 2D), with periodic boundary conditions imposed on its faces.

Starting from the center of the unit cell, each spherical LNP moves according to the Langevin equation subject to a Brownian force F^B^, a Stokes drag force, and a pairwise interaction potential U between the particle and the mucins:

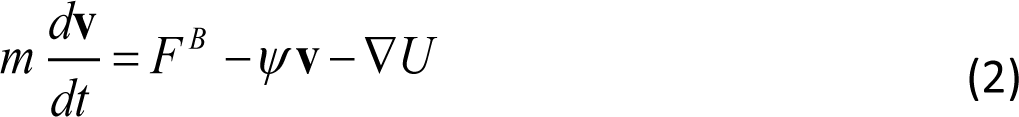

where 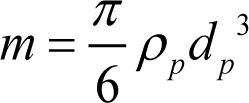 is the mass of particle of density *ρ* and diameter *d*, **v** is its velocity, *t* is the time, Ψ = 3πμ*_f_d_P_* is the drag coefficient and ∇ denotes the spatial gradient. Although inertia is unimportant to NP transport, we retain the particle mass here so we can use an existing module in COMSOL Multiphysics (2020) to solve the Langevin equation and advance the particle positions.[98] We have done numerical experimentation to confirm that this term has negligible effect on our BD results, e.g., the mean-square displacements (MSD), in the parameter range of interest. See Figure S2 of Online Supplementary Material for details of validation. In addition, we have disregarded particle-particle and hydrodynamic interactions in our simulations, similarly to previous studies.[27,72,85] In general, particle-particle interactions start to become significant when the NP volume fraction φ exceeds 2%.[99] In our experiments φ is well below 2%.

The NP-mucin potential U is described in Sec. S2 of the Supplemental Materials. The Brownian force is treated as a Wiener process over a small discrete time step Δ*t* [100,101]:

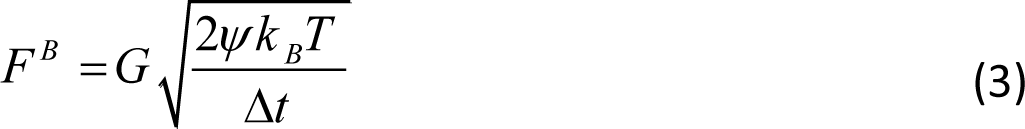

where *k_B_* is the Boltzmann constant, *T* is the absolute temperature, and G is a vector whose components are independent Gaussian random variables of zero mean and unit variance. In all simulations, we set the temperature to 310.15 K, the normal human body temperature. The free diffusivity in the pure solvent (*D_SE_*) follows the Stokes–Einstein relation *D_SE_* = *k _B_Tψ*, relative to which we can quantify the hindrance of particle diffusion hindrance inside the mucus.

### 3.7 Statistical Analysis

Statistical analysis on the experimental data was performed using Prism 9 software (GraphPad). For bar plots, error bars represent standard error of the mean. For Box-and-Whisker plots, the box extends from the 25^th^ to 75^th^ percentiles. The whiskers indicate the entire data range. The statistical significance was determined using one-way ANOVA followed by Tukey or Fisher’s LSD multiple comparison test. A p value ≤ 0.05 was considered statistically significant.

Statistical analysis on the BD simulation data is described in Sec. S3 of the Supplemental Materials.

## Acknowledgment

The authors would like to thank Mitacs Canada (IT19059) and Providence Healthcare (BT, RR, SH, JJF, DS) for the financial support. We acknowledge further financial support from the Deutsche Forschungsgemeinschaft (DFG, German Research Foundation) – Project ID 431232613 – SFB 1449 (SH, DL, SB), and from the NSERC Discovery grant (SH, JF).

