## Supplemental Material for "Exploring Mechanisms of Lipid Nanoparticle-Mucus Interactions in Healthy and Cystic Fibrosis Conditions"

### Supplementary Materials

#### S.1. Size and zeta potential of lipid nanoparticles (LNPs)

The LNP sizes (using dynamic light scattering) and zeta potentials ($\zeta$) were measured using a Zetasizer Nano ZS system (Malvern Instruments Ltd., Malvern, UK) equipped with the Zetasizer software version 7.13 according to standard procedures (Table S1).

**Table S1. Size and zeta potential of mRNA-loaded LNPs**

| PEG concentration | Size $d_p$ (nm) | $\zeta$-potential (mV) |
|---|---|---|
| 1% | 47 ± 19 | - 11 ± 3 |
| 1.5% (baseline) | 42 ± 16 | - 10 ± 5 |
| 2% | 35 ± 10 | - 8 ± 2 |
| 5% | 31 ± 12 | - 12 ± 4 |

In addition, the size of unloaded LNP (1.5% PEG) measured via dynamic light scattering was 38 ± 9 nm. To further confirm the LNP sizes, images of unloaded 1.5% PEG LNPs were captured using cryogenic electron microscopy (Cryo-EM) (Figure S1). The measuring tool in Image J was then used to measure the diameter of unloaded LNPs. The diameter thus obtained, 37.0 ± 5 nm, matches the value obtained by the dynamic light scattering method. For LNP-mRNA sizes, the benchtop LNP + mRNA mixing approach was not feasible for Cryo-EM as it requires very high particle concentrations.

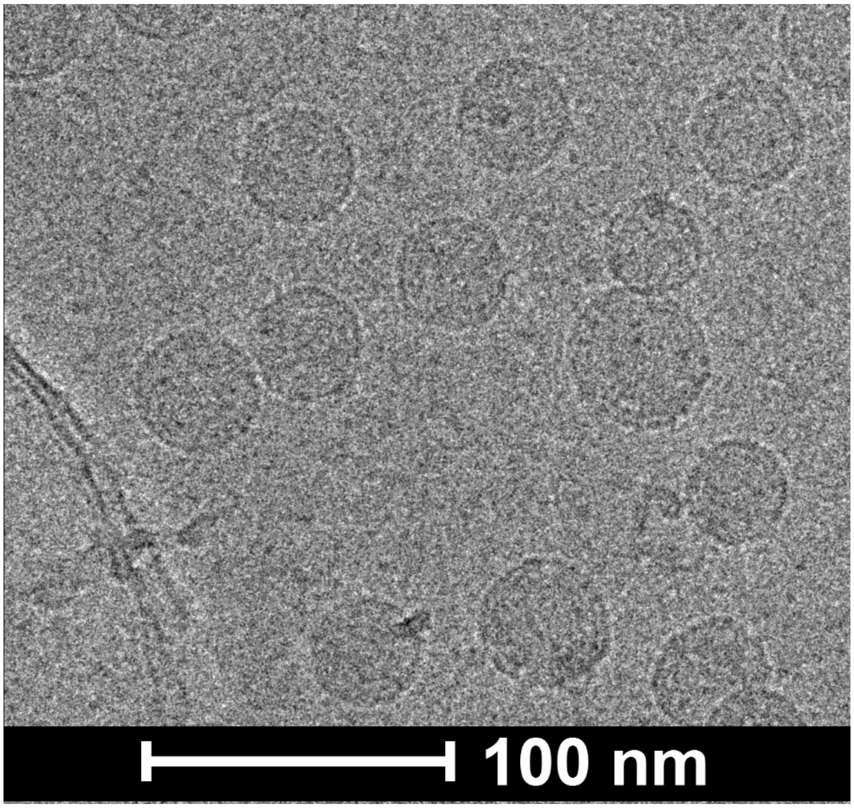


**Figure S1.** A representative image of unloaded LNPs (1.5% PEG) using Cry-EM. To calculate the average size of the unloaded LNP, 143 LNPs were measured as described in section S.1.

### S.2. Interaction potentials

In the Brownian dynamics (BD) simulations, the interaction between the NPs and the mucin polymers is described by a potential U(r) that accounts for steric and electrostatic interactions:

$$U(r) = U^s(r) + U^e(r) \tag{1}$$

Steric interaction: The steric repulsion is modeled by a truncated and shifted Lennard-Jones potential:

$$U^s(r) = \sum_{n=1}^{N} 4\varepsilon \left[ \left( \frac{s}{2r} \right)^{12} - \left( \frac{s}{2r} \right)^{6} + \frac{1}{4} \right] \tag{2}$$

with the cutoff distance of $r_c = 2^{\frac{-5}{6}} s$ (For $r > r_c$, $U^s = 0$). *ε=1 $k_BT$* is the energy depth, *r* the center-to-center distance between the particle and polymer chain, and s=a+$d_p$ is the steric diameter, i.e., the sum of the diameters of the mucin chains, *a =5 nm*[1] and the particle. In addition, *N* is the number of polymer chains in the cubic cell that interact with the NP. To ensure computational efficiency, the summation in Eq. 2 is restricted to a finite number of N neighboring rods. Based on our numerical experimentation, having 12 rods is sufficient for k/b < 0.2, while including 48 rods is necessary for ranges where k/b ≥ 0.2.

Electrostatic interaction: The double-layer interaction between surfaces of various geometries decays exponentially with distance with a characteristic decay length equal to the Debye length.[2] Following Hansing et al.[3], we adopt the following $U^e$ as the potential of NP-mucin electrostatic interaction:

$$U^e(r) = \sum_{n=1}^{N} U_e \exp\left(-\frac{r}{k}\right) \tag{3}$$

where $U_e$ is the strength of electrostatic potential, attractive when $U_e < 0$ and repulsive when $U_e > 0$. Its magnitude should be proportional to the product of the surface charges on the NP and the mucin chains. But it also depends on the geometry of both surfaces, and no generally valid formula is available.[4] Therefore, we have fitted $U_e$ from the diffusivity in a certain baseline case, and then determined its values in other cases by the proportionality to the surface charges. Details are given in Section 2.2 in the main text where NP and mucin surface charges are varied.

The interaction range *k* is the Debye screening length[2]

$$k^2 = \frac{1}{4\pi l_B I} \tag{4}$$

where $I = \frac{1}{2}\sum_j n_j z_j^2$ is the ionic strength and $z_j$ the valence of salt ion *j* and $n_j$ its bulk number density. Besides, $l_B = e^2 / 4\pi\varepsilon_f k_B T$ is the Bjerrum length, *e* the elementary charge and $\varepsilon_f$ the permittivity of the solvent.

### S.3. Statistical Analysis

Our BD simulation tracks the trajectory of *P* particles, each governed by the Langevin equation (see Eq. 2 in the main text). For each of the *P* individual trajectories, we use internal sampling over all pairs of points as a function of the time interval *t*:

$$\overline{q^2(t)} = \frac{1}{M}\sum_{k=1}^{M}\left|q(t_k + t) - q(t_k)\right|^2 \tag{5}$$

where $M$ is the number of all pairs separated by $t$, $t_k = (k\text{-}1)\Delta t$ is the starting time of the $k^{th}$ time step and the starting time of the $k^{th}$ pair, $q$ is the particle position and $\overline{q^2(t)}$ is the single particle MSD. Moreover, we ensemble-average over all the $P$ non-interacting trajectories:

$$\left\langle \overline{q^2(t)} \right\rangle = \frac{1}{P}\sum_{i=1}^{P}\overline{q_i^2(t)} \tag{6}$$

to compute the ensemble MSD $\left\langle \overline{q^2(t)} \right\rangle$ (we will simply call it MSD). Finally, the overall diffusion coefficient, $D$, is calculated:

$$D = \lim_{t\to\infty}\frac{\left\langle \overline{q^2(t)} \right\rangle}{6t} \tag{7}$$

We compared the above method with an ensemble averaging without internal sampling and found that the inclusion of internal sampling effectively reduced the statistical noise by extracting more information from each particle trajectory. In addition, we have validated our numerical scheme against the Stokes-Einstein relation for free-diffusion and a published numerical study (See Figure S2).[3] Details are reported below.

### S.4. Validation of computational model

To ensure accurate BD simulation results in long-time MSD and diffusivity, we have used a sufficiently fine time step $\Delta t$, a sufficiently long running time $T_R$, and a large enough ensemble $P$ for averaging out the stochastic noise Eq. (6). Internal sampling, based on correlations among points on the same trajectory separated by time intervals of different lengths Eq. (5), can help reduce the requirement on the ensemble size $P$. We have carried out detailed numerical experiments to probe how these numerical parameters affect the result, and have determined that the following values offer accurate results at reasonable computational cost:

Dimensionless time step: $\eta = \Delta t\, D_0/b^2 = 10^{-6}$;

Number of particle trajectories: $P = 10^3$;

Total number of time steps: $N_T \geq 10^6$,

Internal samplings: all pairs between two points separated by 1 to $N_T$ - 1 time steps.

These numerical experiments are briefly summarized below.

We calculated the diffusivity $D$ in a pure solvent with $\mu_f$= 0.692 cp, and then compared it with $D_{SE}$ predicted by the Stokes–Einstein relation, with an error defined as $(D-D_{SE})/D_{SE}$. We varied the number of particle trajectories from 250 to 1500 and observed that when P ≥1000, the error is < 5% and decreases over time (Figure S2a). Thus, we set $P$ = 1000. Figure S2(b) shows the effect of refining the time step. For $\eta \leqslant 10^{-6}$, the error is < 5% and decreases over time. Thus we chose $\eta=10^{-6}$ to be the time step. To appreciate the impact of internal sampling on MSD calculation, we compared the errors computed with and without internal sampling in Figure S2(c), which clearly demonstrates how internal sampling reduces the statistical noise in computing the long-time diffusivity $D$.

As further evidence for the appropriateness of these parameter values, Figure S2(d) shows the accurate recapitulation of the Stokes-Einstein relationship for a range of particle size. Figure S2(e) deals with NPs interacting with the mucin chains in our standard setup (see Figure 2D of main text) via steric repulsion and electrostatic repulsion or attraction. Our results show agreement with those of Hansing et al.[5] to within 5%. These results serve as quantitative validations of our BD simulations using the numerical parameters determined in the above.

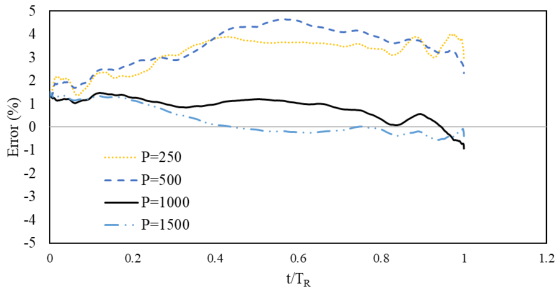


(a)

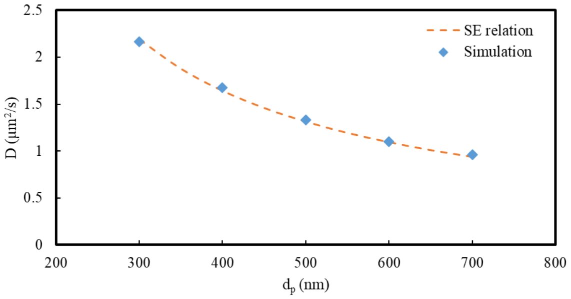


(d)

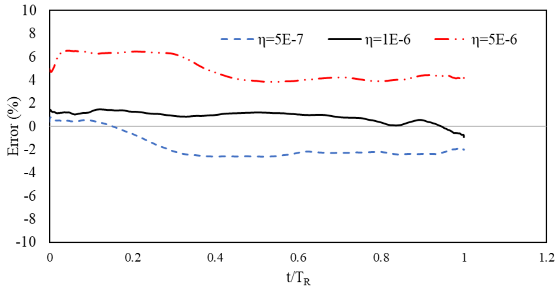


(b)

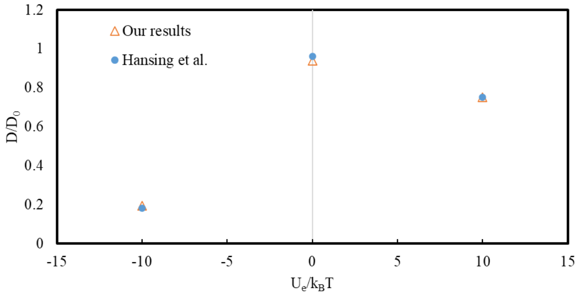


(e)

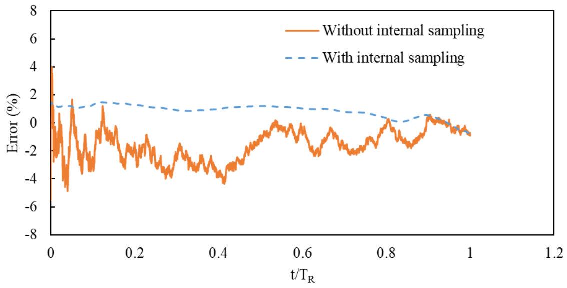


(c)

Figure S2. Error in the diffusivity $(D\text{-}D_{SE})/D_{SE}$ as a function of rescaled time $t/T_R$ for different (a) number of particles (b) time step (c) internal sampling. In all simulations, we set temperature $T$=310.15 K, viscosity $\mu_f$= 0.692 cp, particle size $d_p$=300 nm, and total number of time steps $N_T$ = $10^6$. (d) Comparison between theoretical and numerical diffusivities as a function of NP size. (e) Comparison between our predicted diffusivities with steric and electrostatic interactions and those of Hansing et al.[5] at parameters $\varepsilon/k_BT$=1, $k/b$=0.1, and $s/b$=0.2. A negative $U_e$ represents electrostatic attraction, and a positive one repulsion.
